# The growth-driven penetration strategy of plant roots is mechanically more efficient than pushing

**DOI:** 10.1101/2024.03.16.585213

**Authors:** Yoni Koren, Alessia Perilli, Oren Tchaicheeyan, Ayelet Lesman, Yasmine Meroz

**Author notes:** contributed equally to this work (Yoni Koren, Alessia Perilli).

## Abstract

Plant roots are considered highly efficient soil explorers. As opposed to the push-driven penetration strategy commonly used by many digging organisms, roots penetrate by growing, adding new cells at the tip, and elongating over a well-defined growth zone. However, a comprehensive understanding of the mechanical aspects associated with root penetration is currently lacking. We perform penetration experiments following *Arabidopsis thaliana* roots growing into an agar gel environment, and a needle of similar dimensions pushed into the same agar. We measure and compare the environmental deformations in both cases by following the displacement of fluorescent beads embedded within the gel, combining confocal microscopy and Digital Volume Correlation (DVC) analysis. We find that deformations are generally smaller for the growing roots. To better understand the mechanical differences between the two penetration strategies we develop a computational model informed by experiments. Simulations show that, compared to push-driven penetration, grow-driven penetration reduces frictional forces and mechanical work, with lower propagation of displacements in the surrounding medium. These findings shed light on the complex interaction of plant roots with their environment, providing a quantitative understanding based on a comparative approach.

## 1. Introduction

Plant roots have the extraordinary ability to sense the environment and adaptively grow inside their medium (1), making them one of the most efficient soil explorers amongst living organisms. This is critical for plant survival, enabling water and nutrient uptake, as well as anchoring (2–5). As opposed to the pushing penetration strategy commonly used by many digging organisms, roots penetrate by growing. This unique growth-driven mechanism has recently inspired the development of a new generation of growing robots (6–9).

Root growth generally entails new cells being produced at the tip, just behind the root cap, and an increase in internal turgor pressure, overcoming external resistance, driving their elongation within a fixed sub-apical region called the *growth zone* (2). The cells in the growth zone reach a maximal size, and become part of the *mature zone*, which remains fixed over time, as shown in Fig. 1A. While it is generally accepted that growing roots excel at soil penetration, the underlying mechanism is not well understood. The measured pressure exerted by growing root tips, in the range of 0.1-1 MPa (10–12), is generally lower than the resistance of soil to penetration, in the range of 0.5-4.5 MPa in soft soils (13–15). Traditionally this difference has been attributed to the production of mucilage at the tip of the root which decreases friction (10, 16–20). However, recent observations indicate that soil displacements are highest in the immediate vicinity of the root tip, falling at some distance behind it (19, 21–23), suggesting a relation to the extent of the growth zone. This has motivated work comparing penetration driven by growing versus pushing (18, 24, 25). For example, Sadeghi *et al*. developed a self-growing robot that moves by 3D printing new material at the tip, and demonstrated that the addition of material (growth) facilitated soil penetration, since the rest of the robot (representing the mature zone) is stationary, thus decreasing peripheral friction and energy consumption down to 70% compared to pushing (25). However, there are currently no experimental studies directly comparing the mechanical implications of penetration driven by growth (roots) with penetration driven by pushing.

**Fig. 1.**
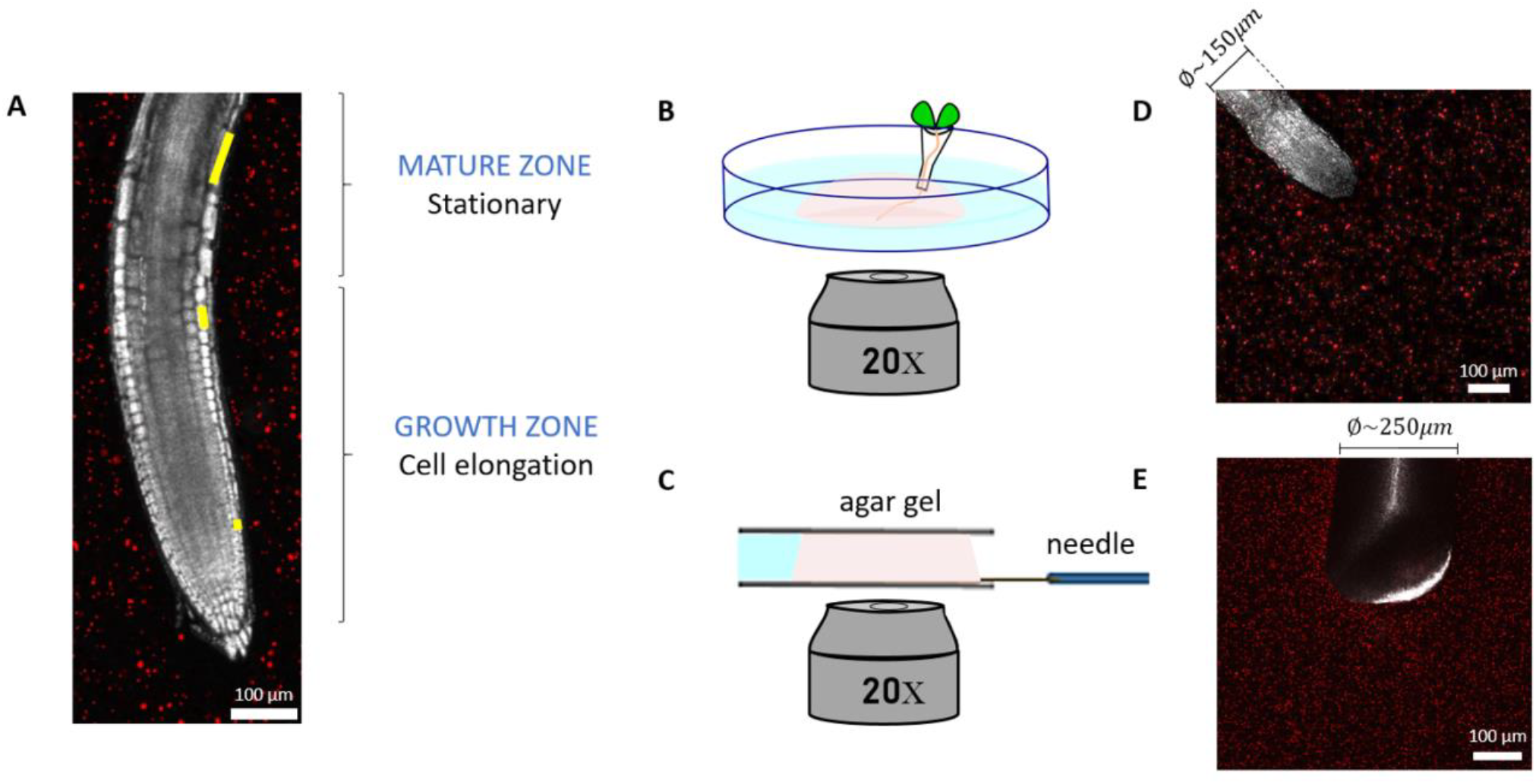
Experimental setup of a growing root and a pushed needle penetrating agar gel. (A) Confocal image of a growing root with its distinct zones: cells elongate within the growth zone, and finally reaching their final size in the static mature zone. Three single cells are marked in yellow at different stages, demonstrating elongation as they get further away from the tip. (B) Schematics showing the glass dish with the Arabidopsis root growing into the agar gel droplet submerged in water and imaged from the bottom using a confocal microscope. (C) Schematics of the needle experiment where the needle is moving horizontally in the agar gel between two cover slips, submerged in water and imaged from the bottom using a confocal microscope. (D, E) Confocal image showing the root tip (D) and needle tip (E) surrounded by agar embedded with fluorescence reporter beads.

While such experimental observations could provide essential insight, the complexity of plant root experiments limits the identification and control of the different contributions, such as friction and changes in mechanical properties of the medium, as well as the role of turgor pressure and cell wall acidification. This motivates the use of experimentally informed computational models, which allow precise control over the problem through the systematic and independent change of relevant parameters. Furthermore, while some aspects of root mechanics and their impact on growth have been elucidated (10, 26–31), and recent theoretical models have described the mechanical interaction of a growing root with an obstacle or its own weight (32– 34), a comprehensive understanding of the mechanical aspects involved in soil penetration of growing roots is still lacking. In particular, computational examinations of object penetration have predominantly concentrated on pushing mechanisms that do not consider growth (17, 20, 35–43), and existing models considering root growth provide limited analyses of the mechanical interaction with the surrounding medium (44–51).

In this work, we present an experimental and computational study comparing growth- and push-driven penetration mechanisms, in terms of mechanical interactions with the surrounding medium. We measure the medium deformation imposed by *Arabidopsis thaliana* roots growing in an agar gel, compared to a pushed micro-needle of similar dimensions, and develop a 3D Finite Elements (FE) numerical model informed by experimental measurements. Based on simulations, we perform a quantitative analysis of these scenarios showing that growing is an energetically favorable penetration mechanism over pushing, leading to smaller deformations in the embedding medium.

## 2 Materials and Methods

### 2.1 Root experiments

Arabidopsis roots were grown in agar hydrogel, and their growing process was recorded in real-time by confocal microscopy. The experimental stages included germination, initial growth, transfer for microscopy observation, and finally tracking the displacements of fluorescent beads embedded in the gel as the root grew further in the gel.

#### 2.1.1 Seed germination

The medium for seed germination was prepared using 0.6% (w/v) Phytagel (Sigma P8169), Murashige and Skoog (MS) salts with vitamins (1.15 gl^-1^) (M0222, Duchefa, 2003 RV Haarlem, Netherlands), MES monohydrate (0.5 gl^-1^) (M1503, Duchefa, 2003 RV Haarlem, Netherlands) and D-(+)-Sucrose (10 gl^-1^) (Bio-Lab Ltd, Israel). Sterile 90mm Petri dishes were filled to a height of approximately 0.8cm with this medium. 0.5-10μl pipette tips (Gilson Clear Micro P10) were filled with the gel in liquid form, using a micropipette, with approximately 5μl of medium from the Petri dish. Once the medium in the tips had solidified, they were cut at approximately 0.5cm from the end and placed upright into the Petri dishes containing solid growth medium; a total of eight to twelve cones were placed in every Petri dish. On top of each of the cones, an Arabidopsis thaliana seed was placed with the help of a sterile wooden toothpick (Supplementary Figure S1). The dish was then sealed and put in a growth chamber to start germination; the growth conditions were 21°C at a 16h light / 8h dark cycle. After four to seven days, shoots grew upwards, and roots grew down the cone towards the aperture.

#### 2.1.2 Preparation of the set-up for microscope imaging

When the roots had grown about 1cm past the cone aperture (between seven to ten days), they were selected for microscope imaging. The roots’ growth was imaged in a second medium, made of 0.6% (w/v) agar gel (Agarose, low gelling temperature; Sigma A9045) supplemented with the same components mentioned in the previously described growth medium. The melted gel was mixed with fluorescent microbeads (FluoSpheres™Carboxylate-Modified Microspheres, 1.0μm, orange fluorescent (540/560), 2% solids; Fischer F8820) in 0.75% concentration (v/v). In order to homogeneously distribute the beads, the mixture was agitated using a vortex mixer for about 1 minute. The cone containing one of the selected roots was lifted from the growth medium using forceps and placed on a glass bottom dish (dish size 35mm, well size 14mm, #1.5 cover glass), with the root lying on the glass and the cone held upright using a piece of PDMS. The gel-beads mixture, still liquid but not too hot so as not to damage the root, was then slowly pipetted on top of the root, forming a hemispherical shaped droplet. Once the gel solidified, the glass bottom dish was filled with water to avoid dehydration during the imaging process. The glass bottom dish was placed in the growth chamber for a few hours to allow the plant to adapt to the new environment.

### 2.2 Needle pushing experiment

The experimental setup for the needle experiment consisted of a needle insertion device (SENSAPEX micromanipulator), the same agar gel which was used for the Arabidopsis thaliana root imaging, and the Zeiss confocal microscope (Zeiss 880, 20X magnification lens). The micromanipulator was placed upon the microscope stage (Supplementary Figure S2), holding and actuating a shaft which contained a stiff tungsten needle of 250μm diameter and length of 1cm, oriented horizontally and located about 100μm above a glass cover slip (#1.5, 24 mm × 60 mm) over the microscope objectives. A 2.5mm-thick strip of silicone rubber was glued to the cover slip in advance, serving as a mold for the agar gel. The vertical distance between the needle and the cover slip was within the working distance of the objective (∼0.55 mm). When the correct position of the needle tip was detected in the microscope, a 1ml agar gel embedded with micro-fluorescence beads (with the same concentration and composition as described in section 2.1.2) was micropipetted and let solidify on the glass coverslip, covering the horizontal needle and surrounded by the silicone strip. This was made to ensure that the front part of the needle was embedded in the gel prior to the start of the experiment. After the gel fully solidified, a second coverslip was placed on top of the gel, holding it gently to prevent any boundary motions of the gel during the needle insertion. Thus, the gel was 2.5mm-thick prior to imaging. The silicone mold also contained a water reservoir keeping the gel moisturized, preventing it from drying. A few drops of water were then carefully added around the gel through a small gap between the upper coverslip and the silicone strip, and the system was then left for stabilization for another 20 min. The needle was gradually inserted into the agar gel using the micromanipulator. The micromanipulator does not allow continuous penetration, and instead we appproximated root growth rate by inserting steps of 2.5μm, at an insertion speed of 100μm/sec (the minimal possible speed for the device). Each step was followed by a 30 min break, to allow the gel to stabilize.

### 2.3 Confocal imaging and agar displacement measurement

For both the needle and the root experiments, the imaging process was performed with an 880 Zeiss confocal microscope using a 20X magnification lens. The 3D root images were captured with an image resolution of 512 × 512 × 100 pixels, with 1.38μm/pixel, for a total area of 708.49 × 708.49 μm^2^ on the xy plane and z-stack of 138 μm. For the root, a time lapse was generated taking Z-stack images at intervals of 4 min for several hours.

Root growth deformed the embedding gel, and displacements were quantified by tracking fluorescent beads embedded in the gel using a Digital Volume Correlation (DVC) algorithm (52–54). Due to instabilities of bead movements at the root-gel interface, displacements around the root were calculated from a minimal distance from the tip (equivalent to 0.6 of the root thickness). The 3D volumetric image was divided into subsets, which were then cross correlated to compute the 3D displacement field of each subset. In our case, the size of the subset was ∼44×44×44μm^3^ and the spatial resolution was 22μm. We measured the typical experimental noise in the system by quantifying the systematic error resulting from imaging artifacts or thermal fluctuations, performing DVC on the same agar gel with no roots/needle imaged over time. We found mean values generally around zero, indicating no global drifts, with fluctuations within 0.3μm and 0.9% for the displacement and strain measurements, respectively (Supplementary Figure S3). These values set the noise threshold for measuring displacements and strains in our system.

For the needle experiments, the 3D confocal image stack was taken after a tip displacement of 2.5μm, and the DVC analysis was performed taking the previous image as a reference. During the 30 min break needed for stabilization period, a z-interval of approximately 300 μm was recorded using the confocal microscope. The image resolution was 700 × 700 pixels, with 1.0 μm/pixel for a total area of 708.49 × 708.49 μm^2^. In order to perform similar analyses in the case of the root and needle, we manually tracked the position of the root tip in all time frames, identifying time points at which the root tip made a 2.5μm displacement. Let *k* be the number of time frames corresponding to such 2.5μm intervals, each *i*-th image was used as a reference for the *(i+k)*-th image in the DVC analysis (for instance, in one case the root penetrated 2.5μm every 4 timeframes, so the first image was correlated to the 5th, the 2nd to the 6th, the 3rd to the 7th and so on). In this way, for each experiment made of *N* images, we obtained a series of *(N-k)* DVC analyses corresponding to the same displacement of 2.5μm and statistically equivalent to each other. In the case of the needle experiments, four DVC analyses were performed for each experiment (one for each time step).

### 2.4 Agar gel characterization

A cylindrical sample of the same agar gel used for the root and needle experiments was prepared using a hollow cylindrical 3D-printed mold with an outer diameter of 40 mm, inner diameter of 14.2 mm and height of 11.4 mm. 1.8 mL of gel was cast inside the mold, which was positioned in the middle of the stage of a DHR-3 rheometer (Discovery Hybrid Rheometer, TA Instruments), between the two loading plates (Supplementary Figure S4a, b). The mold consisted of two parts, so that they could be disassembled after gel polymerization. A thin layer of grease was applied to the inner surfaces of the 3D printed mold. After polymerization and mold removal, the gel was surrounded by a ring-shaped silicone strip, to create a border for a water reservoir keeping the gel moisturized. Rheometer measurements were performed at room temperature and by enclosing the rheometer stage with a polythene sheet to prevent dehydration. Then, the upper plate was manually lowered, until it touched the top surface of the gel evident by non-zero normal forces. The compression test was performed with a slow speed movement of the plate of 0.055 μm/sec, targeting the slow growth movement of roots. Measurements were repeated three times using the rheometer.

The raw data obtained from the compression test of the rheometer at each time point contained the gap distance, which is the distance between the two plates, and the normal force applied by the gel sample on the upper plate. Subsequently, this data was used to calculate the engineering axial strain (ε) and engineering axial stress (σ) of the gel, which are given by:

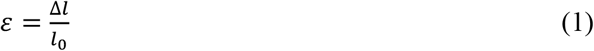

where Δ*l* is equal to the initial height of the sample *l*_0_ minus the gap distance, and

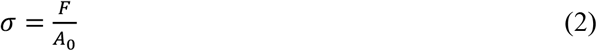

where *F* is the normal force, and A0 is the initial cross-sectional area of the gel sample, that is equal to πd02/4 (d_0_ is the original diameter of the sample). We assumed that the cross-sectional area of the sample did not considerably change throughout the test due to the relatively small change in gap distance. The test data were averaged to extract the resulting stress-strain relation curve (using Eqs. 1 and 2), to be used in the FE model to obtain an elastic-plastic behavior of the simulated agar gel (Supplementary Figure S4c). The elastic regime of the curve was linearly fitted to obtain a Young’s Modulus of 2,361 Pa, a typical value for low-concentrated agar of 0.6% (55, 56). The onset of the plastic regime, which was characterized by approximately constant stress, was measured to be around 5% strain.

### 2.5 Finite Element modeling

For the FE analysis, the ABAQUS Standard/Implicit FE solver (57) in its nonlinear analysis mode was used to simulate the push- and growth-driven penetration of a thin cylindrical and rigid object into a domain, with material properties of the agar gel measured in 2.4. Root parametrization, FE model construction and further details of the simulation setup are detailed in the following sections. To achieve a proper comparison of the push- and growth-driven penetrations, the tip of the penetrating object in both cases (second FE model; see Sec. 2.5.3 for details) was given the same total displacement of 0.52 mm, so that in each increment the tip penetrates the same distance. This way, each of the 100 time points are comparable.

#### 2.5.1 Root growth parametrization

The model of the root geometry was designed based on an experimental image of *Arabidopsis Thaliana* root taken with the confocal microscope. The shape of the root was approximated by a cylinder connected to an elliptical dome, providing a simplified 3D geometry with similar shape and dimensions as the root (Fig. 4A). We assume a vertically positioned root throughout simulations, and disregard gravity effects. To simplify the model system and avoid failure of the material due to extreme deformations, we introduce a burrow within the gel domain, with a diameter corresponding to 0.9 the root diameter; as the root moves in the burrow, it is always in contact and pushes against the burrow wall, leading to friction. To compare the two penetration mechanisms (growing versus pushing) and to account for differences between them, we ensured that the burrow is long enough to allow for sufficient penetration length. Growth occurs only in the growth zone, defined within a fixed region from the tip of length L_gz_. We assume an exponential growth process (49), with a uniform growth rate *ε*_*0*_ (with dimensions 1/sec) within the growth zone. For an infinitesimal segment of length Δ at distance *s* from the tip, the elongation reads:

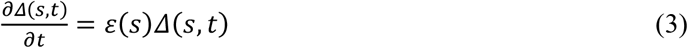

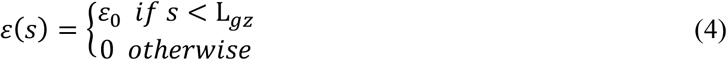

where
In other words, the elongation of the growth zone has the form of exponential growth, where with each time increment dt, Δ is lengthened by *ε*_*0*_Δdt (58). The root body is discretized into vertical sections (*i.e*., elements), which are connected by characteristic key points (generally located at their corners; *i.e*., nodes). Numerically, at every step *n* the length of an element inside the growth zone is updated as follows:

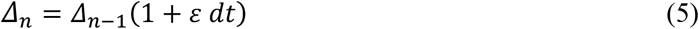

where *dt* is a discretization of time. This procedure determines the evolution of the vertical coordinate (y) of every element. The cross-section coordinates (*x* and *z*) remain unvaried in the cylindrical part of the root, while the convex region (the root tip) is translated as a rigid body (Fig. 4B; see an animated demonstration of the growth in Supplementary Video 1, where for simplicity only the border line of the modeled root is shown). To reduce the computational cost of simulations, we chose a growth zone of L_gz_ = 0.1 mm, smaller than observed values for Arabidopsis ∼1mm (59), without loss of generality. We started with a very dense discretization of the growth zone, to always maintain a significant number of elements inside the growth zone without the need of adding new elements, since elements progressively grow out of the elongation zone and become stationary. In order to avoid high stresses in the elements in the growth zone due to the elongation, ‘surface elements’ were used (57). These elements can support any translation of their nodes without developing stresses, since they have no inherent stiffness. The root geometry was discretized using linear 4-node quadrilateral surface elements, and no material properties were allocated since these elements are computationally considered as rigid compared to the gel.

#### 2.5.2 Model parameters

The material properties of the medium are set based on the rheological measurements detailed in 2.4, with a Young’s Modulus of 2,361 Pa, and the Poisson ratio of 0.49 (37, 60). The plastic region was defined in correspondence to the plateau in the stress-strain curve obtained from the rheological measurements, starting at 5% strain, and the corresponding yield stresses were calculated with the calibration tool in ABAQUS /CAE to match the experimental stress-strain curve (Supplementary Figure S4c). The agar gel was meshed with 8-node linear brick elements, reduced integration and hourglass control, to facilitate the simulation convergence (57). A surface-to-surface contact interaction was defined in the interface between the gel domain and the root’s outer surface, with different values of friction coefficients, to analyze the effect of friction. A static, general step was used to run the simulations, divided into 100 increments, which are referred to as frames, or ‘time points’. Using a static step is reasonable, as the slow processes of root growth and needle insertion can be considered as quasi-static, with the system in equilibrium along the duration of penetration.

#### 2.5.3 Simulations setup and boundary conditions

Two different models were developed. (i) The first model was designed to validate the mechanical response of the computational model by comparing the push-driven simulations and the needle experiment. The penetrating object was initially placed within the modeled agar gel, similar to the starting point in the needle experiment where the gel polymerized around the needle. The diameter and shape of the tip were determined to resemble the actual needle. As for the boundary conditions, the bottom gel surface was constrained in the vertical (y) direction, while the side surfaces were constrained in the lateral (x and z) directions. The penetrating object was constrained in both lateral directions, and was forced to translate 10 μm vertically downwards, similar to the total displacement of the needle (Fig. 3A). We also performed a mesh sensitivity study of the model (see Supplementary Figure S5), finding that further reduction of mesh size did not substantially change the results of FE simulations. However, a finer mesh does significantly increase the computational time. (ii) The second FE model was developed to understand the differences between push- and growth-driven penetrations. Root growth was simulated with distinct zones of elongation and maturation, where the maturation part is stationary and the growth zone elongates to resemble root growth, as detailed in section 2.5.1. Push-driven simulations had similar body geometry but involved rigid-body motion. In both type of simulations, we included a burrow within which the body moves, 10% more narrow than the diameter of the root, to allow access in the front of the penetrator while pushing on the sides during penetration. Similar boundary conditions as in the first model were applied to the agar domain (Fig. 4C).

## 3. Results

### Quantitative characterization of penetration experiments: growing root vs. pushed needle

We perform an experimental characterization of the penetration of *Arabidopsis thaliana* roots growing in an agar gel, and a micro-needle pushed into the same medium (illustrated in Fig. 1B-C). We placed an Arabidopsis root in agar gel and recorded its growth using confocal microscopy at intervals of 4 min during several hours, with an average penetration rate of 10 μm per hour. Similarly, a needle with comparable dimensions was inserted in the same type of gel, performing four consecutive displacements of 2.5μm each, leading to a total penetration of 10μm (see Methods). In order to resolve the 3D displacement field of the gel due to penetration, we follow the displacement of fluorescent beads embedded within the gel (red speckles in Fig. 1A, D-E), and perform a Digital Volume Correlation (DVC) analysis (see Methods). Quiver plots computed by the DVC algorithm for both the root and the needle are shown in Fig. 2A and Fig. 2B, respectively. Assuming radial symmetry, we consider a planar projection of displacements larger than the noise level of the system (∼0.3μm, see Supplementary Figure S3). In both cases, the displacements, mainly detected at the gel interface, were generally oriented in the direction of penetration. We plot the *axial* displacements along a path in front of the tip (illustrated with an orange line in Fig. 2A-B), and the *radial* displacements for two distances from the tip (90μm and 180μm; blue and purple lines in Fig. 2A-B). In order to account for differences in dimensions between the root and needle (the root and needle diameters are considered constant up to the tips and are approximately 150μm and 250μm, respectively, see Fig. 1D-E), we normalize all radial displacements by the penetrator radii (61, 62). Displacements are higher for needle penetration (Fig. 2C-D), suggesting a bigger impact on the surrounding medium. Comparing the radial displacements at 90μm and 180μm from the tip (Fig. 2E-F), we find that while these are similar for the needle penetration, for the growing root displacements are higher closer to the tip. This difference is likely since root elongation (growth) occurs closer to the tip, where it moves relative to the medium, whilst farther away is the mature zone, which remains stationary relative to the medium.

**Fig. 2.**
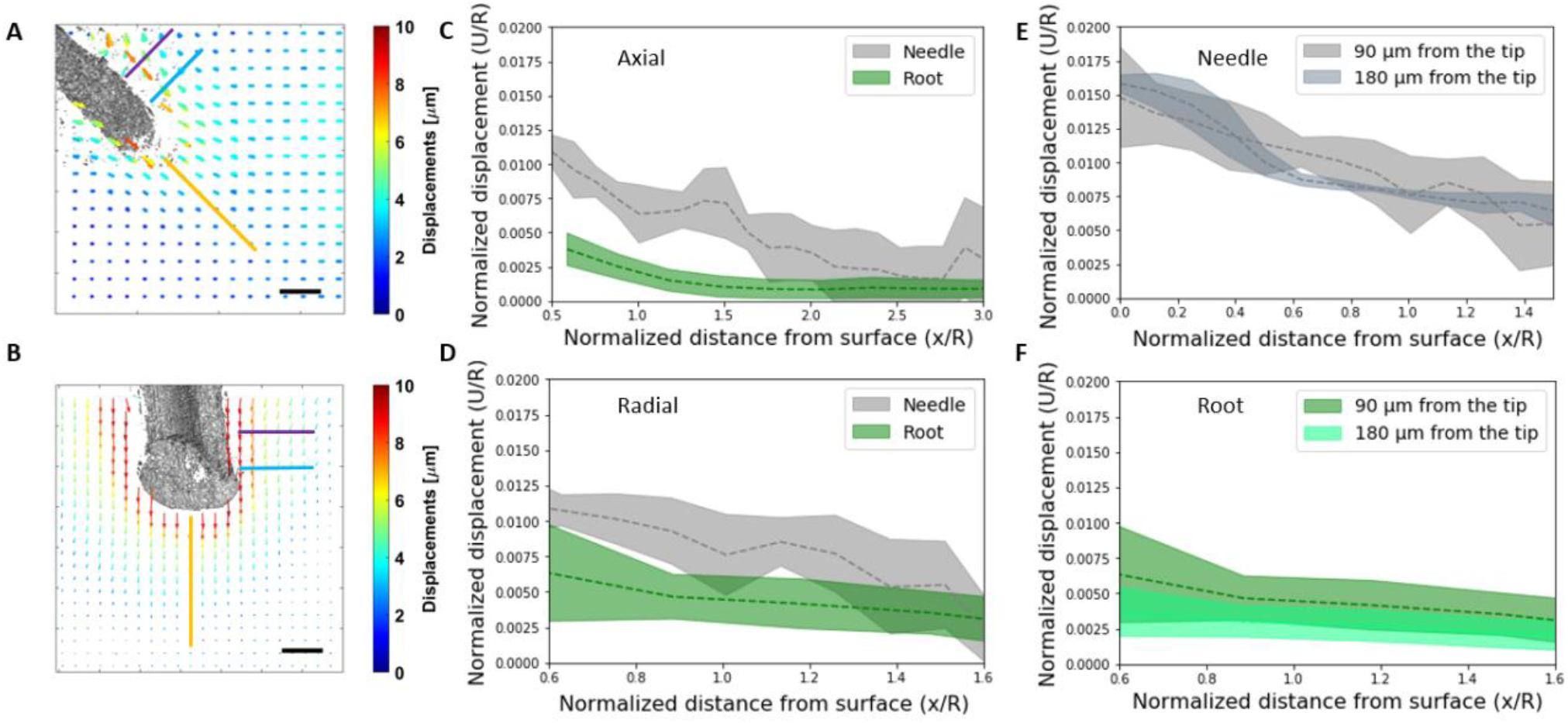
Experimental comparison between a growing root and a pushed needle. (A-B) Examples of displacement fields generated using the DVC approach at a middle cross section of the root and the needle. Color bar and arrow size represents displacement size. Scale bar = 100 μm. Lines represent axial and radial axes for displacement analysis in C-F. (C-D) Comparison of the DVC displacements along: (C) axial axis (orange line in A and B); and (D) radial axis at 90 μm from the tip (blue line in A and B), for a penetration of 2.5 μm. To account for differences in root and needle diameters, 150μm and 250μm respectively, displacements are normalized by the radii. Displacements cannot be measured for points close to the root or needle. The needle leads to greater deformations of the medium. (E-F) A comparison of the DVC displacements along two radial paths, at 90 μm and 180 μm from the tip, as illustrated in A and B (blue and purple lines, respectively). Displacements are similar along the needle, however for the growing root the displacements diminish farther from the tip, suggestive relation to the growth zone.

### Finite Element (FE) computational model for agar gel penetration

In order to obtain a quantitative understanding of the displacement fields observed in the pushing (needle) and growing (root) experiments, we developed a finite element (FE) computational model describing these scenarios (Fig. 3A), described in the Methods. In the simulations, we implemented the geometry of the needle, as well as the mechanical properties of the agar in compression, as measured by a rheometer (see section 2.4 in Methods for further details) and assumed a contact friction coefficient of μ=0.05. To validate that our model captures the mechanical response of the gel, we compared the simulated needle displacements to the measured DVC displacements, shown in Fig. 3B-E for tip penetrations of 2.5 μm and 7.5 μm. We find that the experimental and computational displacement profiles agree along both the axial and radial (90μm from the tip) paths, for both 2.5 μm penetration (Fig. 3F-G) and 7.5 μm penetration (Fig. 3H-I), confirming the model reliably represents the behavior and characteristics of the agar gel in these experiments.

**Fig. 3.**
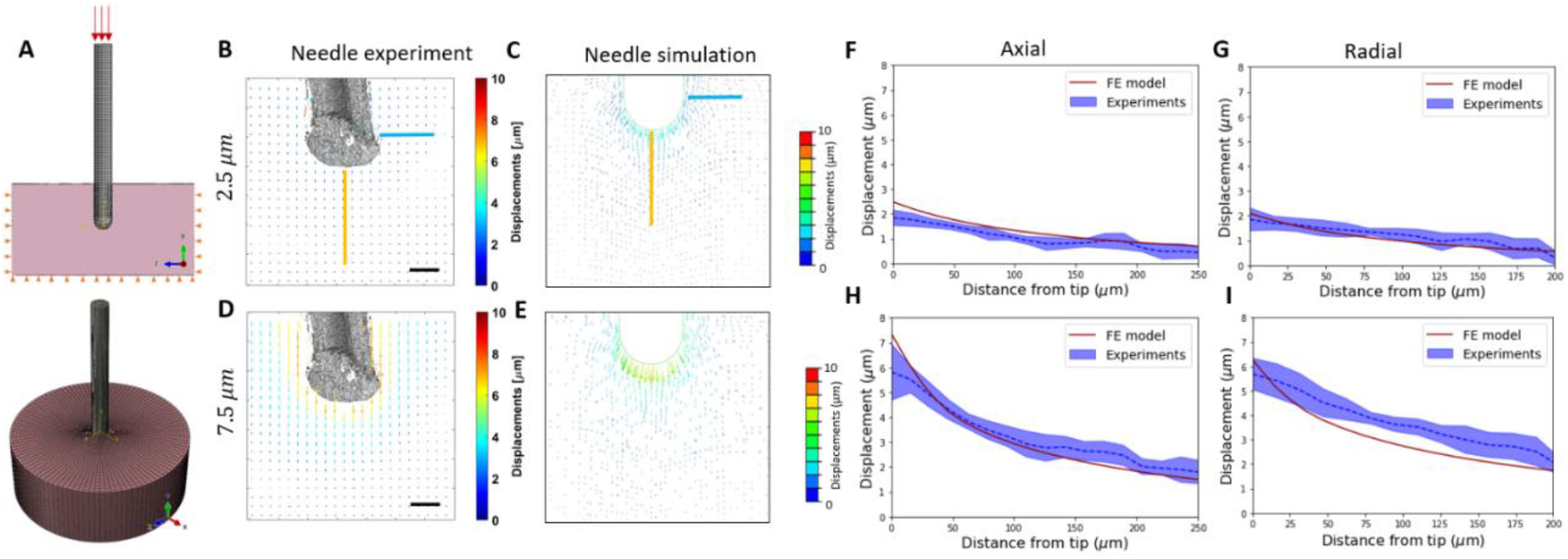
FE Computational model validation. (A) FE modeling of the needle experiment displayed via a middle section view (top) and an isometric view (bottom) of the assembly, including the needle and agar domain. Boundary conditions: orange triangles represent fixed surfaces; red arrows represent the direction of needle displacement. (B-E) Measured and simulated DVC displacement fields at a middle cross section of the pushed needle, following a 2.5 μm penetration (B-C), and following 7.5 μm penetration (D-E). Scale bars = 100 μm. (F-I) Axial and radial displacements (along blue and orange lines in B-C, respectively) at a penetration of 2.5 μm (F, G) and 7.5 μm (H, I). The radial path is 90 μm from the tip. Simulations (red line) and experiments (average curve over 3 experiments in blue dotted line; STD in blue shade).

### Growth reduces frictional forces and mechanical work

Building on our validated numerical framework, we now aim to computationally elucidate the differences between push-driven and growth-driven penetration. We use an identical shape in both types of simulations, informed by *Arabidopsis thaliana* root geometry (Fig. 4A). The simulation of the root growth is based on experimental observations of Arabidopsis roots (47) (see section 2.5.1 in Methods for further details). Fig. 4B shows snapshots of a simulated growing root, where the elongating growth zone is kept at a fixed length Lgz = 0.1 mm (smaller than the observed value L_gz_ ∼ 1 mm in order to reduce computational resources, without loss of generality - see Methods), and the mature zone increases with time. The boundary between the growth and mature zones is marked with a blue line. To capture differences between push-driven and growth-driven penetration, we focus on radial effects due to the dependence on the elongation zone suggested by observations in Fig. 2. To avoid failure of the material due to extreme deformations, we introduce a burrow within the gel domain with a diameter 0.9 of the root diameter; as the root moves in the burrow, it pushes against it, leading to friction (Fig. 4C) – see Methods. We ran simulations for both pushing and growing penetration mechanisms using identical configurations, enabling us to quantitatively study key differences.

**Fig. 4.**
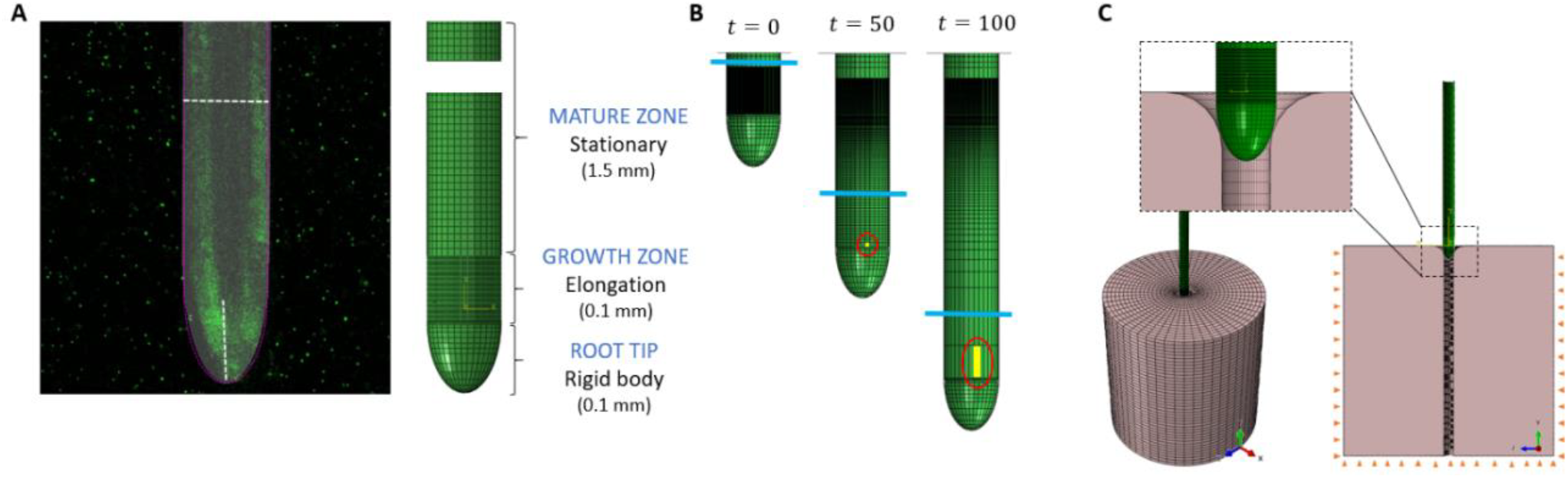
Root growth characterization of the FE model. (A) Confocal image of an Arabidopsis root with measured dimensions of the radius, root tip, and growth zone (left), and the respective FE meshed geometry of the root (right). (B) Snapshots of a growing root simulation. Blue line represents the border between the growth zone of constant length Lgz, and the increasing mature zone. Elements that comprise the growth zone elongate with time, as exemplified by following a single element at the edge of the growth zone marked in yellow. (C) Simulation configuration. An isometric view of the assembly (root and agar domain; bottom left) and a middle section view showing boundary conditions of the agar domain and the burrow (bottom right), with an inset zooming-in on the tip of the root at the beginning of the simulation.

Fig. 5A compares the contact shear force, F_s_, developed at the interface of the root body and the agar for both push and growth simulations following a tip displacement of 0.52 mm, with a friction coefficient μ=0.2. The evolution of forces over time can be seen in Supplementary Movie 2. While the contact shear force is approximately constant along the pushed body, the growing root is characterized by a sharp decrease in forces farther away from the growing tip, where the mature zone stops moving relative to the agar gel (marked by a blue line). These findings hold for other values of μ in the range of 0.05-0.3 (see Supplementary Figure S6). In order to follow the evolution of differences in contact forces with time, *i.e*. with increasing tip displacement, we define the total frictional force (parallel to the direction of penetration) during penetration by summing all the forces acting on the body surface: *F*^*tot*^*(t)*=Σ*i Fs,i(t)*, where *i* denotes the number of a node located on the body surface. The evolution of total frictional force is shown in Fig. 5B. As expected, in the beginning of penetration the force increases with time in an approximately identical manner for both push and growth simulations, as the front part of the body was pushed similarly in both mechanisms. Once the tip penetrates beyond 0.2 mm into the burrow, the static mature zone begins to form, revealing the emerging distinctions in penetration mechanisms. At this point, pushing exhibits higher total frictional force compared to the growing counterpart, for any friction coefficient.

**Fig. 5.**
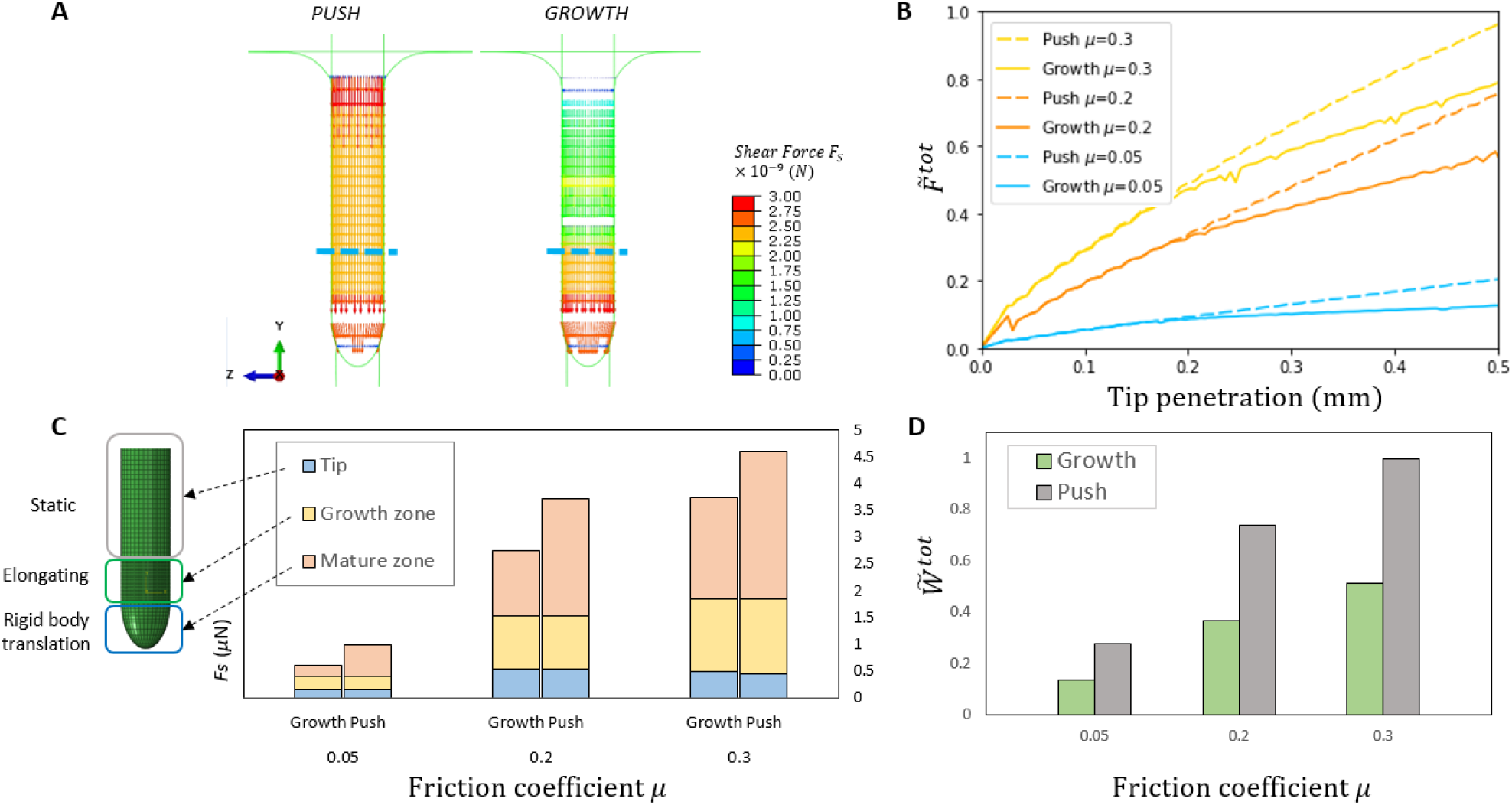
Simulations of push vs. growth penetration in agar. (A) Color maps of frictional shear force Fs at the interface of the root body and the agar following a tip displacement of 0.52 mm) with μ=0.2. While for pushing Fs is constant along the body, for growth-driven penetration shows a clear decrease in the mature zone (the border between the two zones is marked with a blue dotted line; also indicated on the pushed body for comparison). (B) Total frictional shear force Fs^tot^ in the interface as a function of tip displacement. For comparison, values are normalized by the value of Fs^tot^ for pushing with μ=0.3 at tip displacement of 0.52 mm. (C) Local frictional shear force Fs^region^, computed for three regions along the interface: mature zone, growth zone, and tip (referring to the equivalent regions in the pushing simulations where no elongation occurs), following a tip displacement of 0.52 mm, for several friction values. (D) The normalized total mechanical work 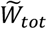 needed for growth is approximately half that required for pushing across multiple friction values (tip displacement of 0.52 mm). Values are normalized by the value of *W*_*tot*_ for the pushed organ with μ=0.3.

Next, to better understand the differences in frictional forces between growing and pushing penetration, we compare the local contact shear forces within different regions along the body - the mature zone, the growth zone, and the tip (Fig. 5C). In the pushing simulation no elongation occurs, and the regions refer to the equivalent positions along the organ. Namely, we measure 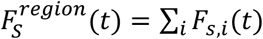 for both the push and the growth simulations, where *i* is the number of nodes located within a specific part of the body surface; see schematics in Fig. 5C. The main differences are evident within the stationary mature zone in the growing simulation and its equivalent region in the pushing simulation, where the pushed organ consistently experiences higher shear forces compared to the growing organ (gray columns). We also compare the mechanical work required for pushing and growing, by measuring the sum of forces acting on the body surface nodes multiplied by the distances each node traveled during penetration: *W*^*tot*^ =Σ*i Fs,i* ⋅ *Δyi*. We find that growing demands approximately half the amount of work in comparison to pushing, across all friction values, suggesting that growing is energetically more efficient than pushing into a medium (Fig. 5D). Increased friction at the interface increases the mechanical work for both growth and push penetration mechanisms, with a constant ratio.

### Growing through the medium decreases the impact on the surrounding medium

To study the extent of the impact of growing vs pushing on the surrounding environment, we compute the propagation of displacements in the medium induced by the pushing/growing object. Fig. 6A-B shows color maps of displacements for μ=0.2, which extend farther around the pushed object compared to the growing one. This is evident in cross-sections of both the top view (Fig. 6A and Supplementary Movie 3) and side view (Fig. 6B, and Supplementary Movie 4). These findings also hold for μ=0.3, with minor differences observed for the lowest friction, μ=0.05 (see Supplementary Figure S7). To quantitatively address the evolution of these differences over time, we focused on two fixed rectangular regions in the agar domain, one 0.1 mm from the top (Fig. 6C), the second at 0.46 mm from the top (Fig. 6D), both 0.14 mm thick (about 1/4 of the total tip displacement) and 0.955 mm long (the distance to the edge of the medium; see Supplementary Figure S8 for more details). At each time point, we compute the total displacements within these regions *U*^*region*^*(t)*=∑*i Ui(t)*for both penetration mechanisms, where *i* is a node located within the region, for different friction levels, normalizing displacements by the final total displacement for a pushed organ with μ=0.3 (Fig. 6C-D). Actual values with no normalization are brought in the insets. As expected, similar displacements are observed in both regions for tip penetration of up to 0.2 mm. However, for larger penetration values, when the mature zone begins to form, pushing leads to higher agar displacements. This difference is particularly pronounced in the vicinity of the mature zone, where displacements plateau for the growing mechanism, in contrast to the pushing mechanism where displacements continue to increase (Fig. 6C). Within the growth zone region, differences between push and growth are less pronounced (Fig. 6D). The raw plots, with no normalization (Fig. 6C-D, insets) show that at a certain tip penetration (∼0.3-0.4 mm; depending on the friction) displacements around the growth zone exceed those around the mature zone, i.e. the front part of the penetrating body has a more pronounced impact on the surrounding medium as penetration increases. In general, higher friction in the root/agar interface increases the differences between pushing and growing.

**Fig. 6.**
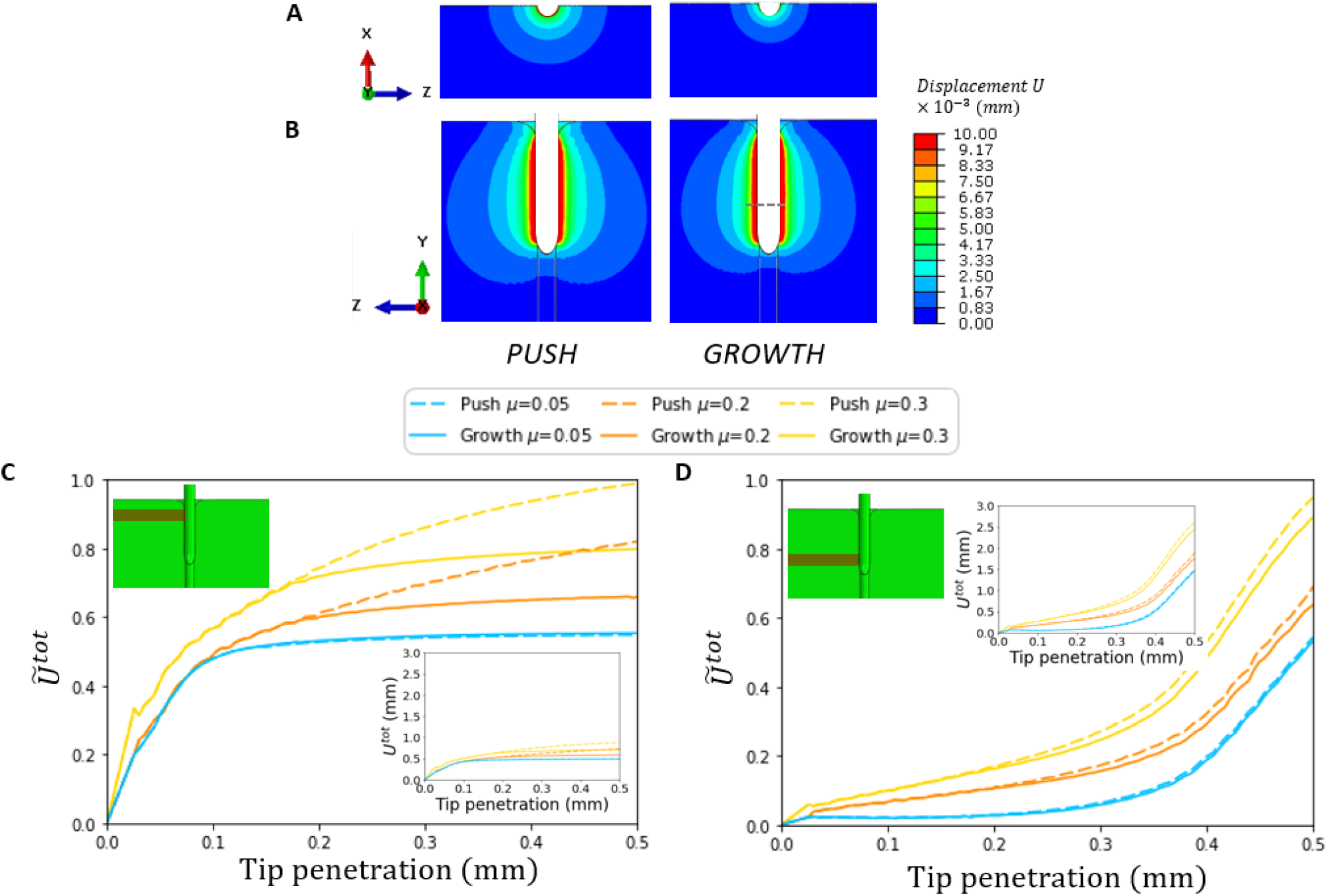
Range of medium deformation due to push vs. growth. (A) Top view and (B) side middle section view, color maps of displacements in the agar domain for pushing and growing, at tip penetration of 0.52 mm, with μ=0.2. The separation between the mature zone and growth zone is marked with a dotted line on the growing body. (C-D) Total displacement Ũ *tot* in two fixed rectangular regions, as a function of tip displacement in the agar domain, both for growing and pushing strategies. Regions are 0.14 mm thick (about 1/4 of the total tip displacement) and 0.955 mm long (the distance to the edge of the medium). (C) The top region is 0.1 mm from the top, shown in the inset relative to the final position of the organ. The tip penetrates past it, such that much of the time the region is in contact with the mature zone. Differences between growing and pushing become evident once the tip penetrates past this region, and it is in contact with the mature zone (or the respective region in the pushed counterpart). (D) The bottom region is 0.46 mm from the top, and most of the time is not in contact with the tip. Only small differences occur between growing and pushing, particularly when the growth zone is in contact with the region. Values are normalized by the total displacement in each region for the pushed organ with μ=0.3 at the final tip displacement. Insets show the absolute displacement values U_tot_.

## 4 Discussion

Here, we studied benefits of the growth-driven penetration mechanism employed by plant roots, compared to commonly used push-driven mechanisms, in terms of mechanical interactions and deformation of the surrounding medium. We measured and compared the displacements within an agar gel induced by the penetration of a growing Arabidopsis thaliana root, as well as a pushed needle. To measure the displacement fields in the gel, we adopted a novel approach (19, 22, 23, 63, 64), combined confocal microscopy imaging and DVC analysis (52–54) of micro-scale deformations in the medium. Experiments revealed that the pushed needle leads to larger deformations in the agar compared to the growing root (Fig. 2).

Motivated by these results, we developed a FE model in order to identify the mechanisms underpinning these differences. We ran simulations, informed by experimental values, and found that growth-driven penetration reduces frictional forces and mechanical work, and causes lower displacements in the surrounding environment compared to the push-driven mechanism. These findings are in line with Sadeghi *et al*. who showed that a self-growing robot, based on additive manufacturing, penetrated granular substrates using up to 50% less force compared to pushing it (24). Mechanical interactions are due to friction, and therefore the movement of the body relative to its surrounding medium. While in the case of a pushed rod the whole body moves against its surroundings, in the case of a growing roots only the tip and sup-apical growth zone move, and the static mature zone does not move. Therefore, new deformations occur only at the front part of the root, and case to develop as the growth zone becomes mature, and therefore stationary. Conversely, when an object is pushed, forces arise from the continuous movement of the entire object, intensifying frictional forces with the domain walls (37). As a result, agar displacements stemming from the pushed object are more substantial and propagate over a greater distance compared to those produced by a growing object.

We note that here we employed a minimal model which captures the dominant characteristics of the problem. However, future studies will explore more intricate material models, such as modeling failure by crack propagation or material rupture (11, 36, 42, 65), characterizing the gel as a viscoelastic fluid adopting a Coupled Eulerian-Lagrangian (CEL)-based FE method for dynamic analysis during the insertion of an object into the medium (37). We also adopted a minimal model of root growth, and future work may include further biological factors which may promote penetration, e.g. lateral roots and root hairs behind the growth zone, which provide anchorage, and mucilage secretion or sloughing cells at the cap, which lubricate the root surface and decrease friction at the root-soil interface, (7, 19, 25, 51).

In all, we find that not only is a growing object energetically more efficient than its pushed counterpart, but that it also leads to a smaller range of deformations, thereby minimizing the environmental disruption it causes to its surroundings. These findings may impact several fields. In the context of thigmotropism, the root’s ability to actively respond to contact and overcome physical barriers (10, 32, 64, 65), this deformation range can be thought of as effectively increasing a root’s presence in the medium, perhaps allowing a neighboring root to sense it through medium deformation – before direct physical contact. Moreover, our findings may provide bio-mimetic solutions for the development of a new generation of growth-driven penetrating devices as an alternative to the traditional push-driven devices, both for soil penetration, as well as for surgical procedures. Indeed, a common goal of many medical intervention surgeries that use catheters, endoscopic devices or biopsy needles is to penetrate complex, curved paths while attempting to minimally damage the surrounding tissue (36, 37), e.g. neurosurgical scenarios (66).

