## Supplementary material for "The growth-driven penetration strategy of plant roots is mechanically more efficient than pushing"

**Supplementary Videos:**

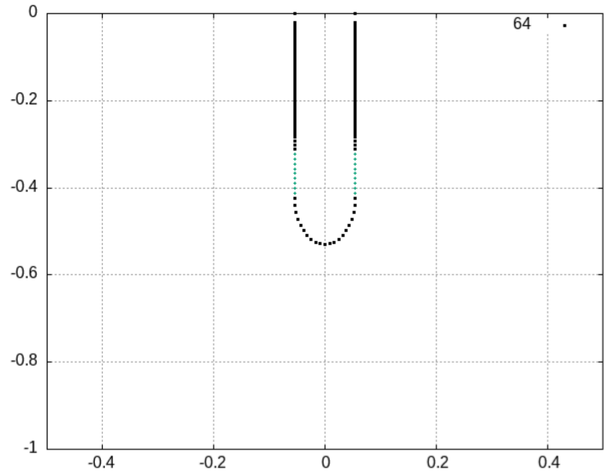


**Supplementary Video 1.** Growth animation in MATLAB. For simplicity, only the border line of the modeled root is shown. The green dots are mobile, while the black dots are stationary at each time point of the simulation to achieve the ‘extension from the tip’ characterization of the root.

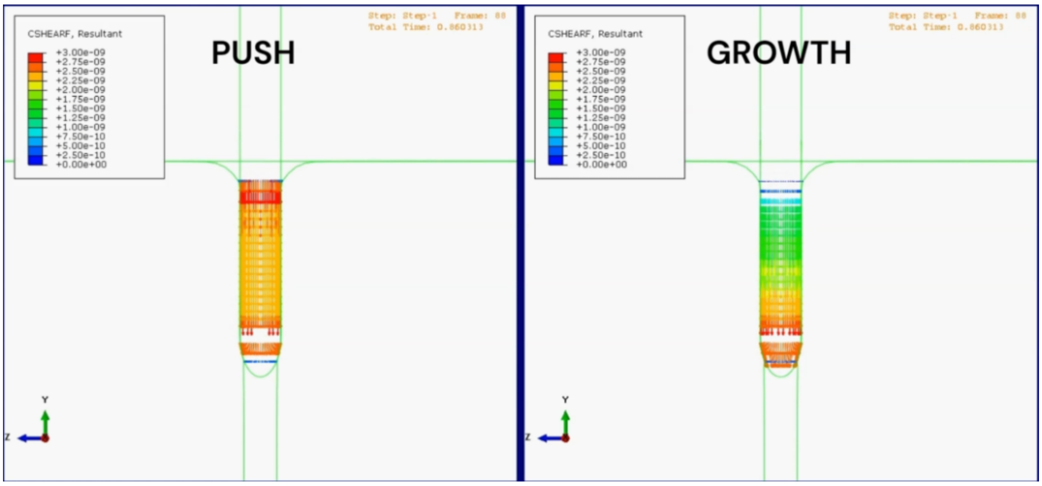


**Supplementary Video 2.** Contact shear forces $F_s$ developed at the interface of the root body and the agar throughout the push and growth simulations (left panel and right panel, respectively), with a friction coefficient μ = 0.2. The forces are measured in Newton (N).

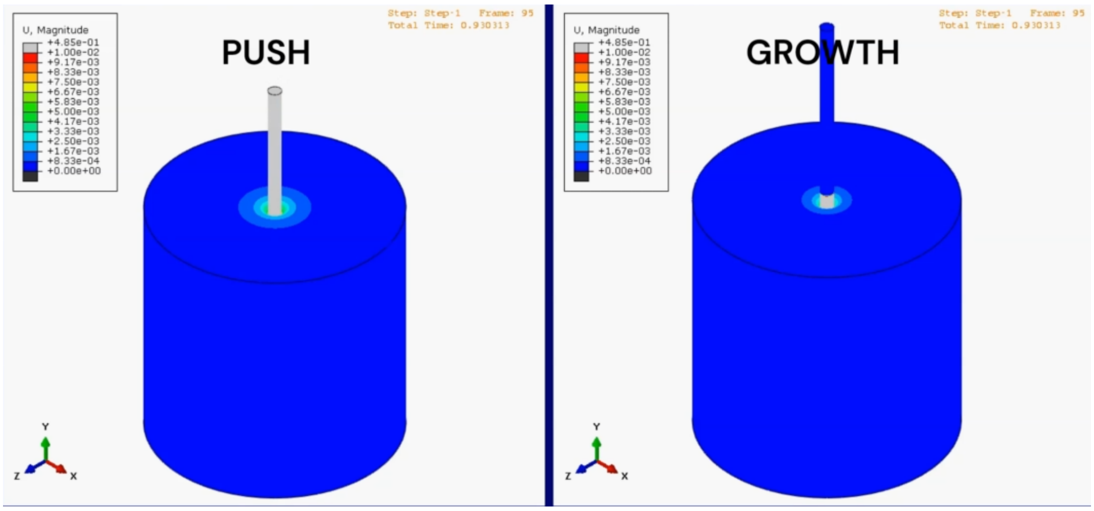


**Supplementary Video 3.** Displacements propagating in the agar due to the penetration of the organ throughout the push and growth simulations (left panel and right panel, respectively), with a friction coefficient μ = 0.2, top view. Displacements are measured in millimeters (mm).

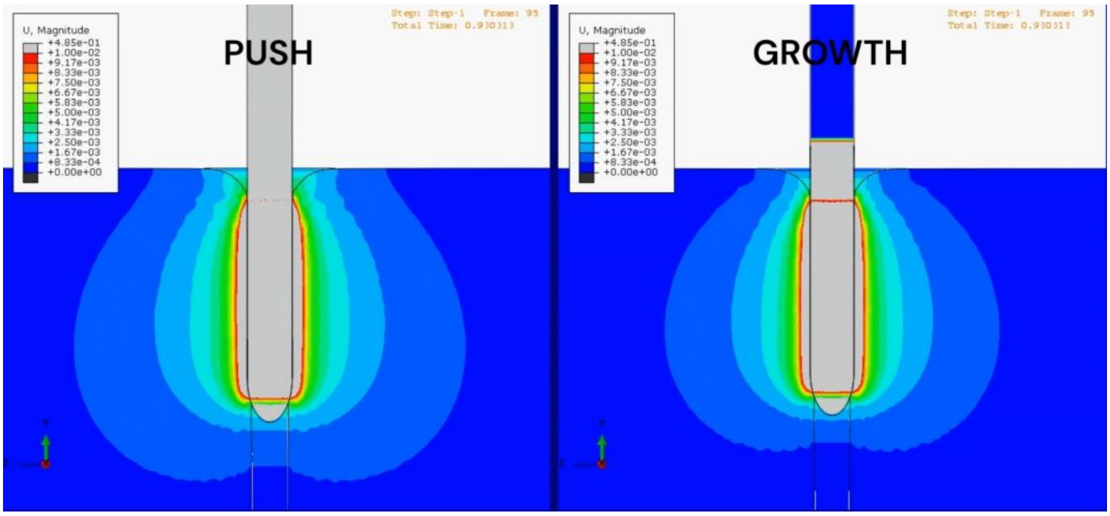


**Supplementary Video 4.** Displacements propagating in the agar due to the penetration of the organ throughout the push and growth simulations (left panel and right panel, respectively), with a friction coefficient μ = 0.2, side middle section view. Displacements are measured in millimeters (mm).

### Supplementary Figures:

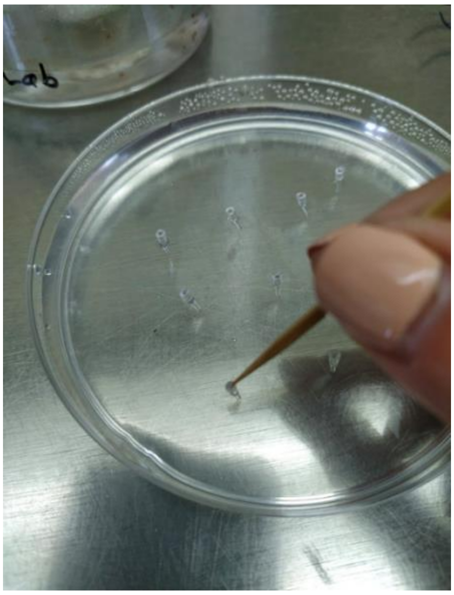

**Fig. S1.** An example of a Petri dish with cones inserted in the agar gel and the seeds being placed on top of them with the help of a wooden toothpick.

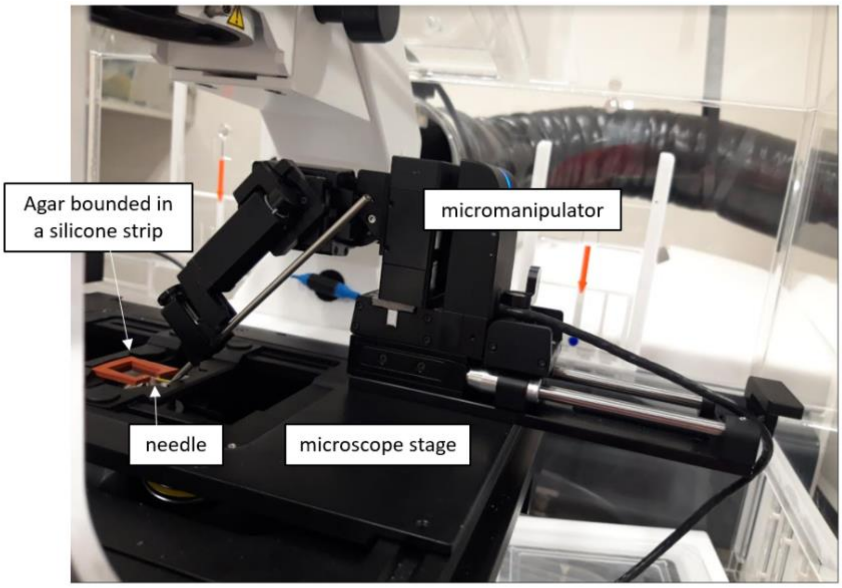


**Fig. S2.** Experimental setup for the needle experiment. A micromanipulator device with a shaft that holds the needle, to be inserted into an agar gel bounded by a silicon rubber strip (orange). The system is placed upon the Zeiss confocal microscope, to image the needle insertion into the gel.

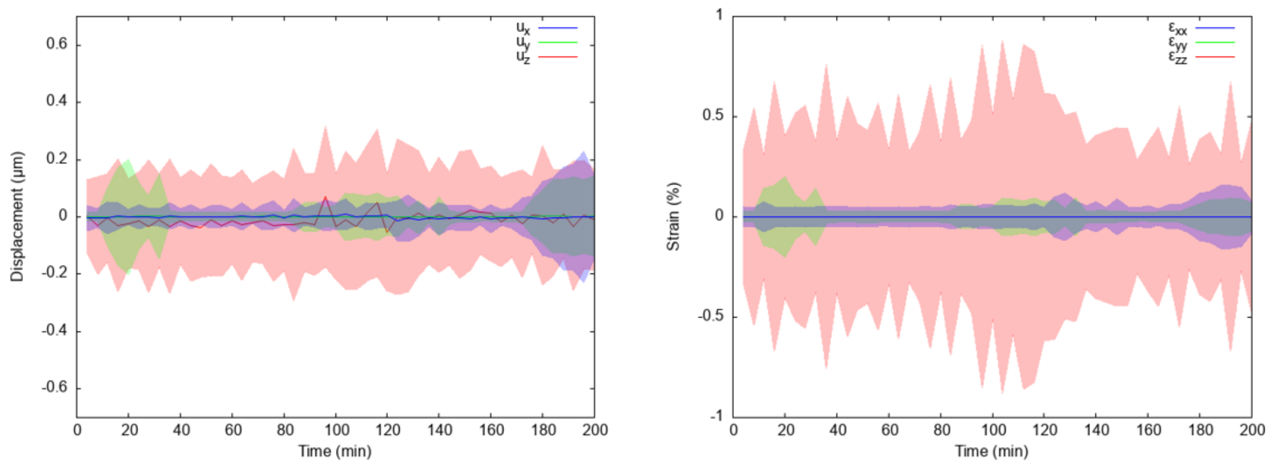


**Fig. S3.** DVC control for the roots and needle experiments. x, y and z components of average displacement (left panel) and strain (right panel) as a function of time for an agar gel without root/needle. The averages are correctly ~zero, while the standard deviation (within ±0.3μm for displacements and ±0.9% for strains) gives an indication of the noise level of the system, due to numerical error and thermal fluctuations of the gel.

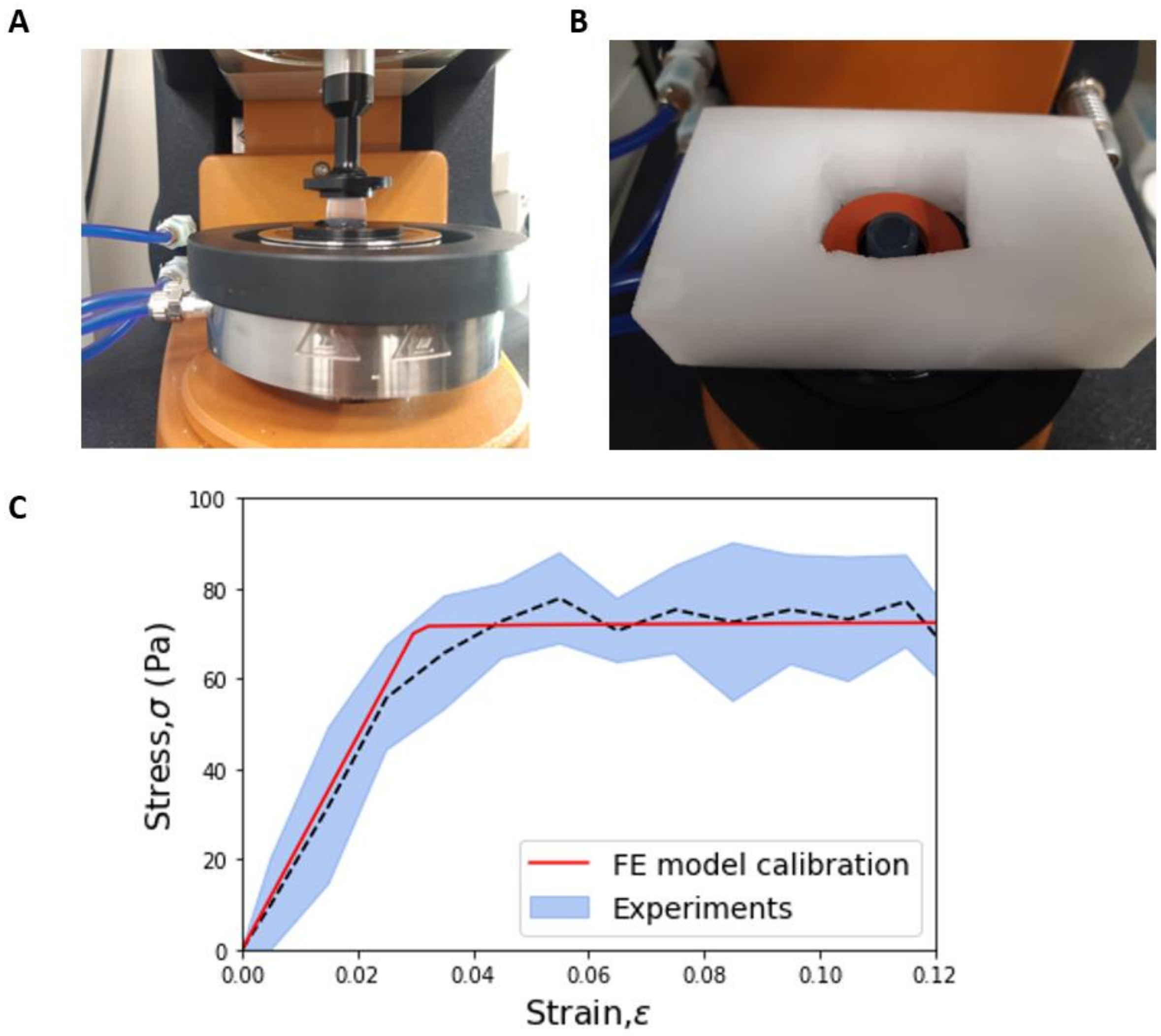


**Fig. S4.** Agar mechanical parameters characterization. (A) Demonstration of gel compression test using the DHR-3 rheometer. (B) The gel sample surrounded by water confined with silicon strip and a water saturated sponge. (C) Compression stress-strain response. Dotted line indicates the measured stress (data was averaged over 3 experiments; STD in blue shade), whereas the red solid line indicates the fitted stress-strain curve approximation, which defines the gel mechanical behavior used in the FE model.

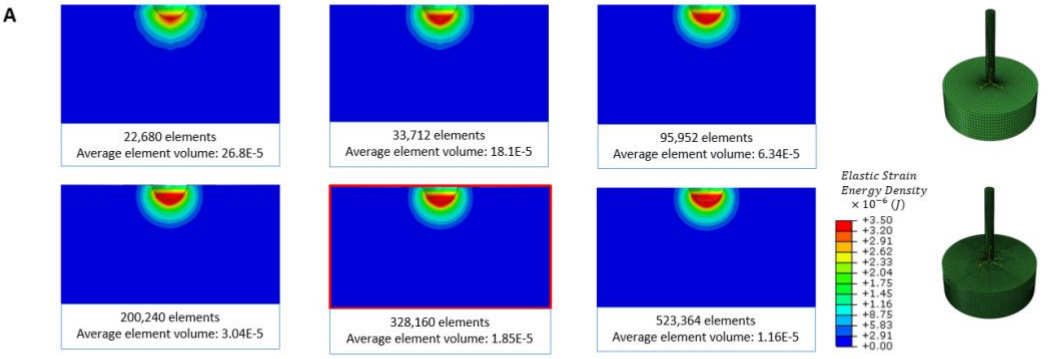


B

| Number of mesh elements | Elastic energy [J] | Relative error (%) |
|---|---|---|
| 22,680 | 5.42E-08 | 2.50 |
| 33,712 | 5.35E-08 | 1.00 |
| 95,952 | 5.31E-08 | 0.27 |
| 200,240 | 5.29E-08 | 0.01 |
| 328,160 | 5.30E-08 | 0.14 |
| 523,364 | 5.29E-08 | - |

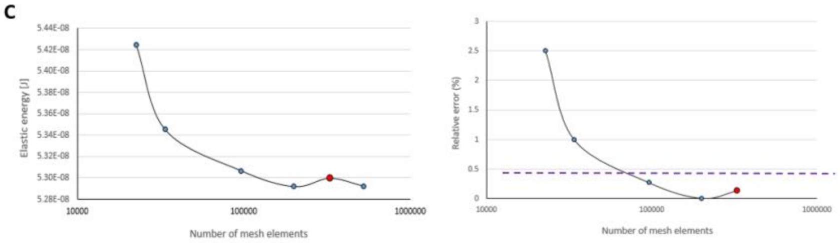


**Fig. S5.** Mesh sensitivity study. (A) Middle section view color plot of the elastic strain energy density in the elements of the agar domain around the needle tip for the push validation model; friction coefficient of the object/agar interface of 0.05. The mesh that was chosen for the validation mentioned in the paper is marked in red. Right: typical coarse and fine meshes. (B) Comparison of the six meshes tested – the one that was chosen for the validation mentioned in the paper (written in red) along with the other five meshes. Values of the total elastic energy in the agar domain elements are reported, as well as the relative error between the total elastic energy calculated for each mesh and the finest mesh (appearing in the sixth row). (C) The total elastic energy in the agar domain elements as a function of the number of mesh elements (left), and the error in the total elastic energy in the agar domain elements relative to the finest mesh, as a function of the number of mesh elements (right). The mesh that was chosen for the validation mentioned in the paper is marked in red. A relative error below 0.5% is considered sufficiently low. Further reduction of mesh size hardly affected the results; however, it significantly increased the computational time.

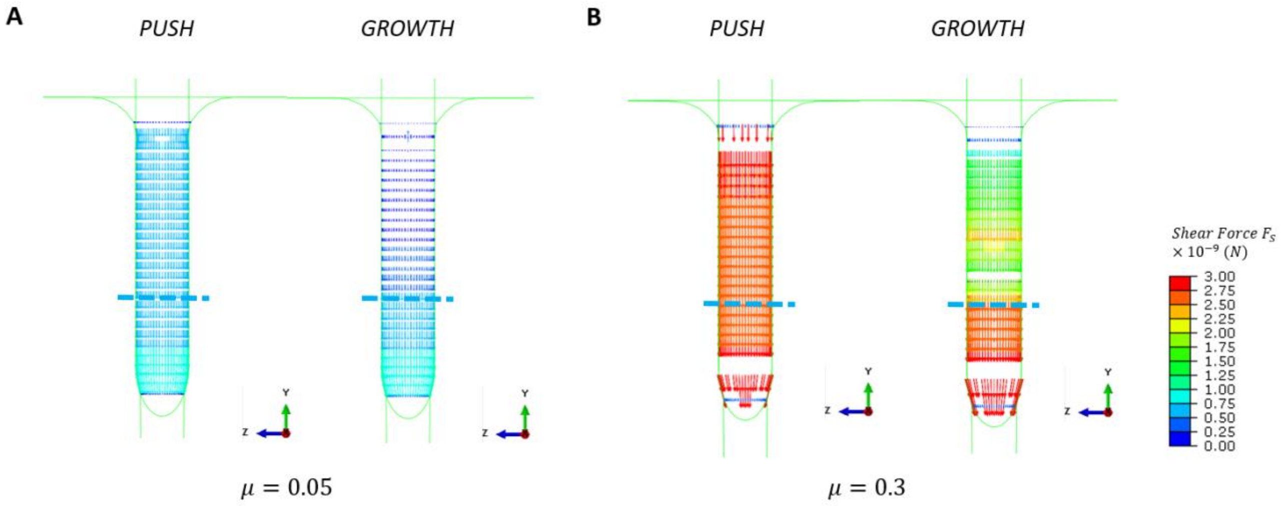


**Fig. S6.** Simulations of push versus growth penetration into an agar burrow: middle section view color maps of frictional shear force $F_S$ at the last time point (tip displacement of 0.52 mm), for: (A) μ=0.05; and (B) μ=0.3. The separation between the growth zone and the mature zone is marked by a blue dotted line.

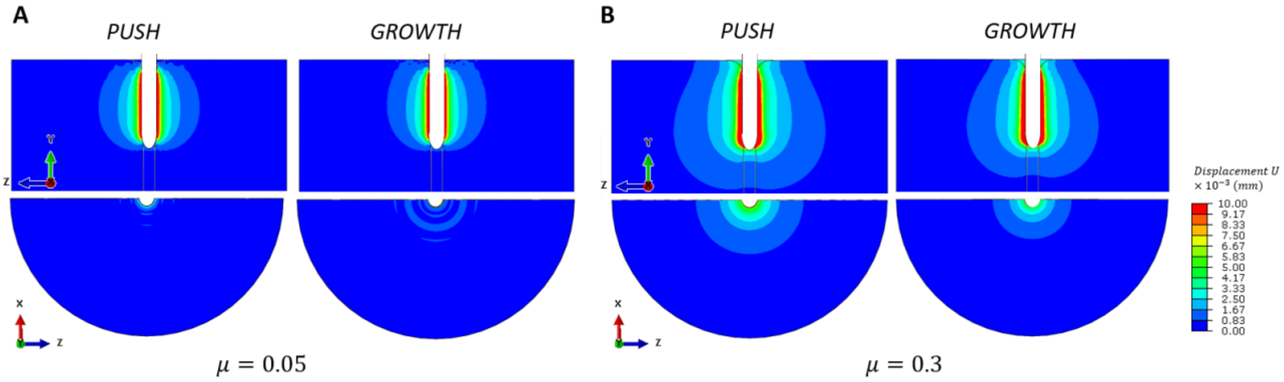


**Fig. S7.** The effect on the agar domain in push versus growth simulations: displacements propagation in the agar domain. (A) Middle section view and top section view color maps of the displacements for each of the two penetration mechanisms at the last time point (tip displacement of 0.52 mm), for (A) μ=0.05; and (B) μ=0.3.

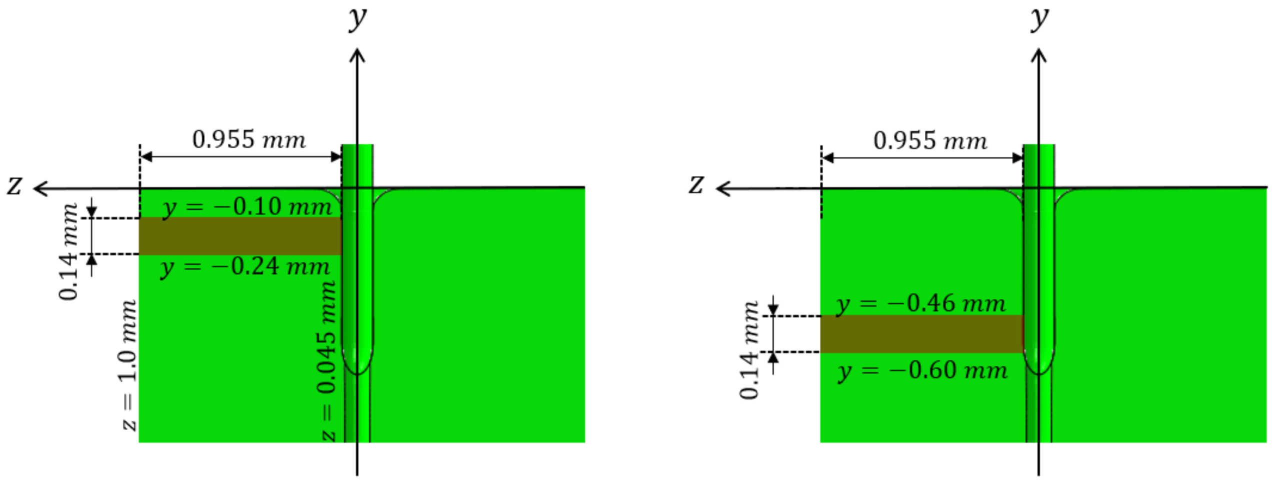


**Fig. S8.** The locations and dimensions of the two fixed rectangular regions in the agar domain, where the total displacements were calculated. Left: corresponds to Fig. 6C; right: corresponds to Fig. 6D.
